# MONI-*k*: An index for efficient pangenome-to-pangenome comparison

**DOI:** 10.1101/2022.08.09.503358

**Authors:** Travis Gagie

## Abstract

Maximal exact matches (MEMs) are widely used in bioinformatics, originally for genome-to-genome comparison but especially for DNA alignment ever since Li (2013) presented BWA-MEM. Building on work by Bannai, Gagie and I (2018) and again targeting alignment, Rossi et al. (2022) recently built an index called MONI that is based on the run-length compressed Burrows-Wheeler Transform and can find MEMs efficiently with respect to pangenomes.

In this paper we define *k*-MEMs to be maximal substrings of a pattern that each occur exactly at least *k* times in a text (so a MEM is a 1-MEM) and briefly explain why computing *k*-MEMs could be useful for pangenome-to-pangenome comparison. We then show that, when *k* is given at construction time, MONI can easily be extended to find *k*-MEMs efficiently as well.

## 1 Introduction

A maximal exact match (MEM) of a pattern *P* [1..*m*] with respect to a text *T* [1..*n*] is a substring *P* [*i*..*j*] of *P* that occurs in *T* but such that either *i* = 1 or *P* [*i −* 1..*j*] does not occur in *T*, and either *j* = *n* or *P* [*i*..*j* + 1] does not occur in *T*.^1^ MEMs were originally used to compare genomes [10] but became especially popular among bioinformaticians after Li [11] used them for DNA alignment in BWA-MEM.

Because BWA-MEM and other well-known approaches — such as suffix trees and fully bi-directional FM-indexes, which can add and delete characters at either end of the pattern; see [12, 14] — cannot handle more than a few genomes efficiently,^2^ Bannai, Gagie and I [3] described a version of the r-index [7] that finds all MEMs of *P* with respect to *T* and can handle pangenomes efficiently.^3^ Their data structure occupies *O*(*r*) words of space by itself, where *r* is the number of runs in the Burrows-Wheeler Transform (BWT) of *T*, and requires two passes over *P* and an auxiliary data structure supporting fast random access to *T*.^4^ If a single random access takes *f* (*n*) time, then their data structure takes *O*(*mf* (*n*)) time.

Rossi et al. [16, 17] (see also [4, 9]) implemented Bannai et al.’s data structure in an index called MONI, eventually targeting alignment. They used a balanced straight-line program (SLP) for *T* to support random access in *O*(log *n*) time,^5^ so MONI finds all MEMs of *P* with respect to *T* in *O*(*m* log *n*) time. In a separate paper [5], they and their coauthors observed that if the SLP is used to support longest common extension (LCE) queries in *O*(log^2^ *n*) time, then MONI needs only one pass over *P* and *O*(*m* log^2^ *n*) time. They named their one-pass implementation PHONI because of the increased running time, but later realized that if the SLP is locally consistent as well as balanced then the LCE queries take *O*(log *n*) time.

Ahmed et al. [2] modified PHONI to obtain software for recognizing DNA strands to eject from nanopore sequencers, which they called SPUMONI. Their initial idea was that, rather than finding MEMs, SPUMONI should compute online the matching statistics [6] of a strand with respect to a pangenome of the target species, and ejects the strand if the distribution of lengths in those matching statistics seems too much like the distribution of lengths in the matching statistics between unrelated DNA samples. They later found that computing the exact lengths in the matching statistics was unnecessary, and numbers they called pseudo-lengths could be used instead and computed more quickly (and, in fact, gave better results).

In this paper we consider a generalization of MEMs, which we call *k*-MEMs. A *k*-MEM of *P* with respect to *T* is a substring *P* [*i*..*j*] of *P* that occurs at least *k* times in *T* but such that either *i* = 1 or *P* [*i* − 1..*j*] does occurs fewer than *k* times in *T*, and either *j* = *n* or *P* [*i*..*j* + 1] occurs fewer than *k* times in *T*. This is a generalization of MEMs because a MEM is a 1-MEM and vice versa. Figure 1 shows an example. The latest implementation of MONI can find all the *k*-MEMs of *P* with respect to *T* in *O*(*m*(log *n* + *k* log log *n*)) time, since it can compute matching statistics and enumerate matches occurrences in *O*(log log *n*) time per occurrence — but the linear dependence on *k* is unappealing.

**Fig. 1.**
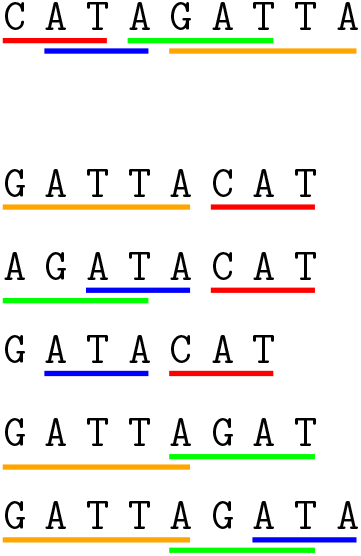
The 3-MEMs of CATAGATTA with respect to the set of strings GATTACAT, AGATACAT, `GATACAT`, GATTAGAT, GATTAGATA. Notice `TAGAT` occurs both in CATAGATTA and in the set of strings — but only twice in the set, so it is not a 3-MEM. If we were to insert CATAGATTA itself into the set, however, then `TAGAT` would be a 3-MEM and `AGAT` would not (because it would no longer be maximal); the other 3-MEMs would remain 3-MEMs (since they are allowed to occur more than 3 times).

To see why computing *k*-MEMs could be useful in pangenome-to-pangenome comparison, suppose we have two pangenomes, *T*_1_ consisting of 100 genomes from people with symptoms of a genetic illness and *T*_2_ consisting of 1000 genomes from people drawn from the general population (the pangenomes need not be disjoint). If we can compute for a genome *G* all the 20-MEMs of *G* with respect to *T*_1_ and all the 5-MEMs of *G* with respect to *T*_2_ — that is, perform a genome-to-pangenome comparison — then we can identify substrings of *G* that are fairly common in *T*_1_ (occurring at least 20 times in 100 genomes) but fairly rare in *T*_2_ (occurring fewer than 5 times in 1000 genomes): that is, any substring of *G* contained in some 20-MEMs of *G* with respect to *T*_1_ but not contained in any 5-MEM of *G* with respect to *T*_2_.^6^ If we do this for each of the 100 genomes in *T*_1_, then we may gain insight into what substrings are more typical of genomes of people with the illness. The ability to choose *k >* 1 is important here, because substrings typical of recessive illnessses or ones with multifactorial causes are likely to appear in both pangenomes at least once, in which case plain MEM-finding is not informative. Of course, being able to choose *k >* 1 also lets us compare a genome *G* in *T*_1_ to all of *T*_1_ meaningfully; since it is in *T*_1_, the whole of *G* (or, at least, each chromosome) is a 1-MEM with respect to *T*_1_.

In Section 2 we show how, for any given *k*, MONI can easily be extended to find all the *k*-MEMs of *P* with respect to *T* in *O*(*m* log *n*) time while still occupying *O*(*r* + *g*) words of space, where *r* is again the number of runs in the BWT of *T* and *g* is the number of rules in a given balanced and locally-consistent SLP for *T*. We call this extension MONI-*k*. We note that with our approach, *k* must be given at construction time; we leave as future work developing a similar data structure that takes *k* at query time. In the near future, we plan to extend the implementation of MONI to obtain an actual implementation of MONI-*k*, and apply it.

Because Section 2 mainly consists of a fairly technical case analysis, in Section 3 we give a simplified example of how MONI-*k* works, finding the MEMs of CATAGATTA with respect to GATTACAT, AGATACAT, `GATACAT`, GATTAGAT, GATTAGATA.

## 2 Extending MONI to MONI-*k*

The simplest version of MONI consists a run-length compressed BWT of the text *T* [1..*n*] with suffix-array (SA) entries stored for each position *i* at a run boundary — that is, *i* = 1, *i* = *n*, BWT[*i*] ≠ BWT[*i −* 1] or BWT[*i*] ≠ BWT[*i* + 1] — and a balanced, locally consistent SLP for *T*. This occupies *O*(*r* + *g*) words of space, where *r* is the number of runs in the BWT and *g* is the number of rules in the SLP.

Consider a character BWT[*i*] and a subsequence BWT[*j*_1_], …, BWT[*j*_*k*_] in the BWT of *T* such that BWT[*j*_1_], …, BWT[*j*_*k*_] are copies of BWT[*i*] and the only ones in BWT[*j*_1_..*j*_*k*_]. Notice BWT[*i*] can be included in BWT[*j*_1_], …, BWT[*j*_*k*_]. If there is no such subsequence BWT[*j*_1_], …, BWT[*j*_*k*_] then there are fewer than *k* copies of BWT[*i*] in *T*, so no *k*-MEM can contain a copy of BWT[*i*].

If there is such a subsequence, then we can find it quickly by computing LF(*i*) (that is, finding the position in the BWT of the character that precedes BWT[*i*] in *T*); finding an interval of size *k* including LF(*i*) such that the minimum LCP value in that interval is maximized (by starting with the interval [LF(*i*)] and greedily decrementing the top endpoint of the interval or incrementing the bottom endpoint, depending on which LCP value is greater, until the interval has size *k*); and then evaluating LF^*−*1^ on the endpoints of the interval. We note as an aside that with this construction, if we want to change *k* then we can rebuild the index without rebuilding the BWT — but we leave explaining the details of the construction to the full version of this paper, which we will prepare when we have MONI-*k* fully implemented and tested.

Let *ℓ* be the length of the longest common prefix of the suffixes *T* [SA[*i*]*−*1..*n*], *T* [SA[*j*_1_] − 1..*n*] and *T* [SA[*j*_*k*_] − 1]..*n*] of *T*. We say BWT[*j*_1_], …, BWT[*j*_*k*_] are a *close k-subsequence* for BWT[*i*] if *T* contains strictly fewer than *k* occurrences of *T* [SA[*i*] *−* 1..SA[*i*] *−* 1 + *ℓ*]. In other words, the suffixes of *T* starting at BWT[*j*_1_], …, BWT[*j*_*k*_] have the longest common prefix with the suffix starting at BWT[*i*] of any set of *k* suffixes of *T*.

For each character BWT[*i*] at a run boundary in the BWT, we choose a close *k*-subsequence BWT[*j*_1_], …, BWT[*j*_*k*_] for BWT[*i*] and store *j*_1_, SA[*j*_1_], *j*_*k*_ and SA[*j*_*k*_]. This adds *O*(*r*) more words to MONI’s space usage; note we store only the endpoints of the subsequence, even if there are gaps in the subsequence. If *k* = 1 then the close *k*-subsequence for BWT[*i*] is BWT[*i*] itself, so we need store only SA[*i*]; this is what we did in previous versions of MONI, which found 1-MEMs.

Suppose we are searching for the *k*-MEMs of *P* [1..*m*] with respect to *T* and, after we have processed *P* [*q*..*m*], we have an interval BWT[*s*_*q*_..*e*_*q*_] of length at least *k* such that the suffixes *T* [SA[*s*_*q*_]..*n*], *T* [SA[*s*_*q*_ + 1]..*n*], …, *T* [SA[*e*_*q*_]..*n*] of *T* have the longest common prefix with *P* [*q*..*m*] of any set of *k* suffixes of *T*. Furthermore, suppose we know SA[*s*_*q*_], SA[*e*_*q*_] and the length *ℓ*_*q*_ of that longest common prefix. Notice we are not assuming BWT[*s*_*q*_..*e*_*q*_] is maximal: for example, *T* [SA[*e*_*q*_ + 1]..*n*] might have an equally long common prefix with *P* [*q*..*m*], or an even longer one (although in that case that prefix must occur fewer than *k* times in *T*). We consider three cases and prove the same lemma in each case:

### Case 1

If every character in BWT[*s*_*q*_..*e*_*q*_] is equal to *P* [*q −* 1], then we can perform a standard backward step and obtain an interval BWT[*s*_*q−*1_..*e*_*q−*1_] of length at least *k* such that the suffixes

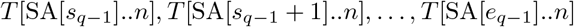

of *T* have the longest common prefix with *P* [*q −* 1..*m*] of any set of *k* suffixes of *T*.

**Lemma 1 (for Case 1).** *The suffixes*

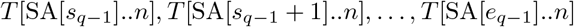

*of T have the longest common prefix with P* [*q −* 1..*m*] *of any set of k suffixes of T*.

*Proof*. Immediate from the definition of the BWT.

Notice SA[*s*_*q−*1_] = SA[*s*_*q*_] *−* 1 and SA[*e*_*q−*1_] = SA[*e*_*q*_] *−* 1 and the longest common of prefix of *T* [SA[*s*_*q−*1_]..*n*], *T* [SA[*s*_*q−*1_ + 1]..*n*], …, *T* [SA[*e*_*q−*1_]..*n*] and *P* [*q −* 1..*m*] has length *ℓ*_*q−*1_ = *ℓ*_*q*_ + 1. In this case we use *O*(log log *n*) time.

### Case 2

If some character in BWT[*s*_*q*_..*e*_*q*_] is equal to *P* [*q −* 1] but some other is not, then some character BWT[*i*] at a run boundary in BWT[*s*_*q*_..*e*_*q*_] is equal to *P* [*q −* 1]. Let BWT[*j*_1_], …, BWT[*j*_*k*_] be the close *k*-subsequence we stored for BWT[*i*]. We forget *s*_*q*_ and *e*_*q*_ and perform a standard backward step from the interval BWT[*j*_1_..*j*_*k*_], which contains exactly *k* copies of *P* [*q −* 1], to obtain an interval BWT[*s*_*q−*1_..*e*_*q−*1_] of length exactly *k*.

**Lemma 1 (for Case 2).** *The suffixes*

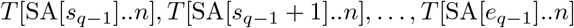

*of T have the longest common prefix with P* [*q −* 1..*m*] *of any set of k suffixes of T*.

*Proof*. Because BWT[*i*] is in BWT[*s*_*q*_..*e*_*q*_], we know *T* [SA[*i*]..*n*]’s longest common prefix with *P* [*q*..*m −* 1] is the longest prefix of *P* [*q*..*m −* 1] that occurs at least *k* times in *T*. Therefore, by the definition of a close *k*-subsequence, *T* [SA[*j*_1_]..*n*], …, *T* [SA[*j*_*k*_]..*n*] have the longest common prefix with *P* [*q*..*m*] of any set of *k* suffixes of *T*.

Since BWT[*j*_1_] = … = BWT[*j*_*k*_] = BWT[*i*] = *P* [*q −* 1], it follows that

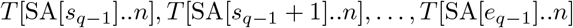

have the longest common prefix with *P* [*q −* 1..*m*] of any set of *k* suffixes of T.

Notice SA[*s*_*q−*1_] = SA[*j*_1_] − 1 and SA[*e*_*q−*1_] = SA[*j*_*k*_], and the longest common of prefix of

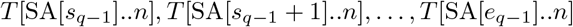

and *P* [*q −* 1..*m*] has length *ℓ*_*q−*1_ = min(LCE(SA[*s*_*q*_], SA[*j*_1_]) + 1, *ℓ*_*q*_ + 1). In this case we use *O*(log log *n*) time.

### Case 3

If no character in BWT[*s*_*q*_..*e*_*q*_] is equal to *P* [*q*], then we find the last copy BWT[*i*] of *P* [*q*] in BWT[1..*s*_*q*_ − 1] and the first copy BWT[*i*^*′*^] of *P* [*q*] in BWT[*e*_*q*_ + 1..*n −* 1]. BWT[*i*] and BWT[*i*^*′*^] must be at run boundaries in the BWT, so we have stored the starting positions *j*_1_ and 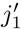 and ending positions *j*_*k*_ and 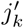 of their close *k*-subsequences, along with 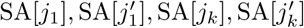. If

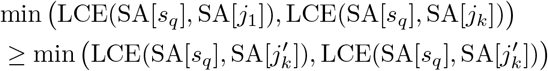

then we forget *s*_*q*_ and *e*_*q*_ and perform a standard backward step from the interval BWT[*j*_1_..*j*_*k*_] to obtain an interval BWT[*s*_*q−*1_..*e*_*q−*1_] of length exactly *k*; otherwise, we perform a standard backward step from the interval 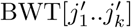 to obtain BWT[*s*_*q−*1_..*e*_*q−*1_]. To simplify our presentation and without loss of generality, assume the inequality above holds (the other situation is symmetric).

**Lemma 1 (for Case 3).** *The suffixes*

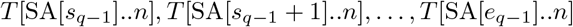

*of T have the longest common prefix with P* [*q −* 1..*m*] *of any set of k suffixes of T*.

*Proof*. Because BWT[*s*_*q*_] is in BWT[*s*_*q*_..*e*_*q*_], we know *T* [SA[*s*_*q*_]..*n*]’s longest common prefix with *P* [*q*..*m*] is the longest prefix of *P* [*q*..*m*] that occurs at least *k* times in *T*. By the definition of the BWT, either *T* [SA[*i*]..*n*] or *T* [SA[*i*^*′*^]..*n*] has the longest common prefix with *T* [SA[*s*_*q*_]..*n*] of any suffix of *T* preceded by a copy of *P* [*q −* 1], but we cannot directly check which because we do not store SA[*i*] and SA[*i*^*′*^].

We can check indirectly by computing the lengths of the longest common prefixes of *T* [SA[*j*_1_]..*n*], …, *T* [SA[*j*_*k*_]..*n*] and *T* [SA[*j*_1_*′*]..*n*], …, *T* [SA[*j*_*k*_*′*]..*n*] with *T* [SA[*s*_*q*_]..*n*], which are

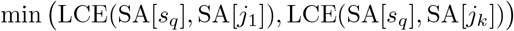

and

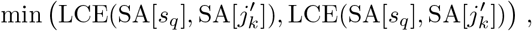

respectively. Recall that we assume the former is at least the latter.

It follows that *T* [SA[*j*_1_]..*n*], …, *T* [SA[*j*_*k*_]..*n*] have the longest common prefix with *P* [*q*..*m*] of any set of *k* suffixes of *T* all preceded by copies of *P* [*q −* 1]. Therefore,

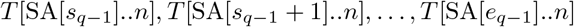

have the longest common prefix with *P* [*q −* 1..*m*] of any set of *k* suffixes of *T*.

Notice that when the inequality above holds, SA[*s*_*q−*1_] = SA[*j*_1_] *−* 1 and 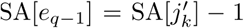 and the longest common prefix with *P* [*q −* 1..*m*] of any set of *k* suffixes of *T* has length

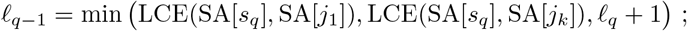

the other situation is symmetric. In this case we use *O*(log *n*) time.

Summarizing our results from all three cases and applying induction, we have the following theorem:

### Theorem 1.

*Suppose we have an interval* BWT[*s*_*q*_..*e*_*q*_] *of length at least k such that the suffixes*

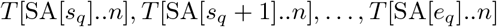

*of T have the longest common prefix with P* [*q*..*m*] *of any set of k suffixes of T*. *Furthermore, suppose we know* SA[*s*_*q*_], SA[*e*_*q*_] *and the length ℓ*_*q*_ *of that longest common prefix. Then in O*(log *n*) *time we can find an interval* BWT[*s*_*q−*1_..*e*_*q−*1_] *of length at least k such that the suffixes*

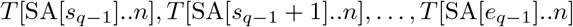

*of T have the longest common prefix with P* [*q −* 1..*m*] *of any set of k suffixes of T*. *Simultaneously, we find* SA[*s*_*q−*1_], SA[*e*_*q−*1_] *and the length ℓ*_*q−*1_ *of that common prefix*.

Whenever *ℓ*_*q−*1_ *≤ ℓ*_*q*_, we know

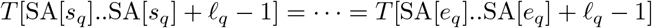

is a *k*-MEM. This gives us the following corollary:

### Corollary 1.

*Given (1) a text T* [1..*n*] *whose BWT consists of r runs, (2) a balanced and locally consistent SLP for T with g rules, and (3) a positive integer k, we can build an instance of MONI occupying O*(*r* + *g*) *words of space such that later, given a pattern P* [1..*m*], *in O*(*m* log *n*) *time we can find the k-MEMs of P with respect to T*. *For each k-MEM we obtain the location of one of its occurrences in T*.

MONI-*k* is the implementation described in Corollary 1. We note that, as with MONI, we can add *O*(*r*) words to MONI-*k* such that, after finding the *k*-MEMs, we can list the occurrences of any of them using constant time per occurrence [7, 15].

## 3 Example

We hope a simplified example of how MONI-*k* works will complement the proofs in Section 2. Suppose *T* is the set of strings GATTACAT, AGATACAT, `GATACAT`, GATTAGAT, GATTAGATA and we want to find 3-MEMs. We build and store a rank data structure over the run-length compressed BWT as we would for regular MONI but, instead of storing the SA samples at run-boundaries in the BWT, for each character at a run boundary we compute and store its close 3-subsequence and the SA samples at the endpoints of its subsequence. Figure 2 shows the BWT — actually the eBWT [13], to simplify the presentation — for our example and Figure 3 shows the ranges between the endpoints of the close 3-subsequences and the lengths of the longest common prefixes associated with those ranges (although we do not store the lengths, which we do not store, since we can compute them with the LCE data structure).^7^ Although it does not happen in our example, we note that a row number need not be an endpoint of the associated close 3-subsequence. We do not show the SA samples nor the LCE data structure, since the key point is how we use the close 3-subsequences.

**Fig. 2.**
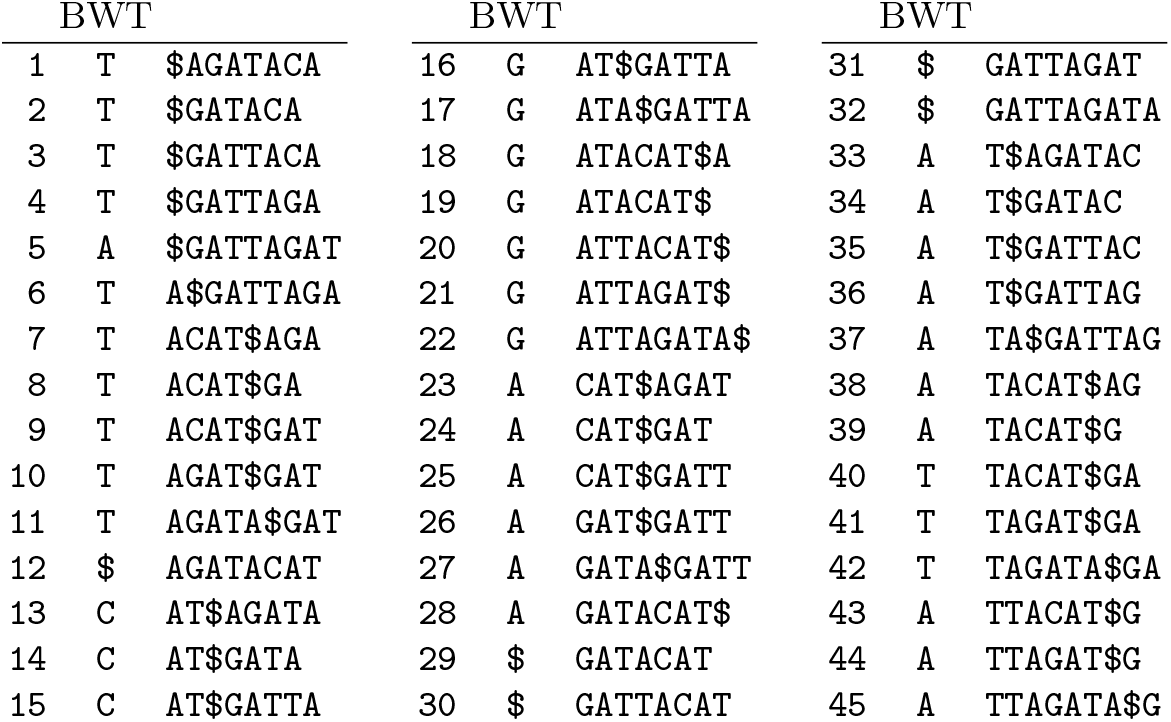
`The eBWT for the set of strings GATTACAT, AGATACAT, GATACAT, GATTAGAT, GATTAGATA.`

**Fig. 3.**
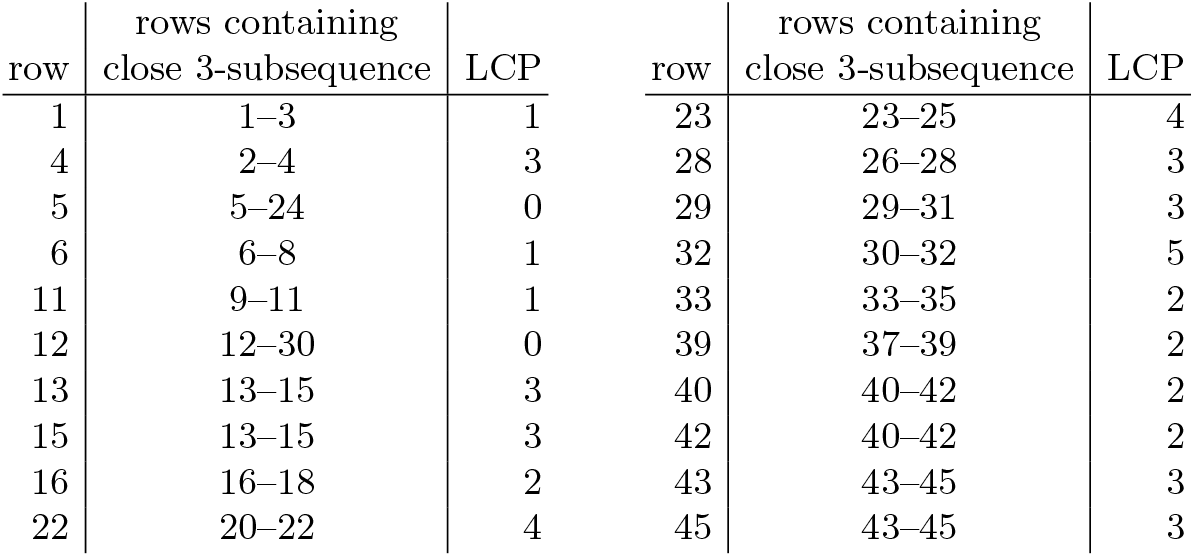
The ranges of the close 3-subsequences for our running example, and the lengths of the longest common prefixes associated with those ranges. Since we store a range for each endpoint of each of the *r* runs in the BWT — *r* = 11 in this case — and an SA entry at the endpoint of each range (not shown), storing all the ranges and the SA sample takes *O*(*r*) space (on top of the space for regular MONI). We show the lengths of the longest common prefixes, but we need not store them since we have the LCE data structure.

To find the 3-MEMs of *P* [1..9] = CATAGATTA with respect to the indexed set *T* of strings, we first choose the close 3-subsequence of an occurrence of *P* [9] = `A` at a run-boundary, say 23–25. This leaves us in Case 1 with a match length of 0 (not including the `A`s), so we simply backward step and find 3 copies of *P* [8] = `T` in rows 7–9. This is again Case 1, now with a match length of 1, so we simply backward step and find 2 copies of `A` and 1 copy of `T` in rows 39–41.

This leaves us in Case 2 with a match length of 2 — since *P* [7] = `T` again — so we change the range to the close 3-subsequence 40–42 of the `T` in row 40, and set the match length to the minimum of the current match length 2 and the length 3 of the longest common prefix of the corresponding suffixes (which we can find by passing the SA samples to the LCE data structure, neither of which are shown in order to keep the figures simple).

We again backward step and find 3 copies of *P* [6] = `A` in rows 43–45. This is again just Case 1, now with a match length of 3, so we backward step and find 3 copies of *P* [5] = `G` in rows 20–22. (Please bear with us: when we reach the left edge of the last 3-MEM `GATTA` in *P*, this will get more interesting.) This is yet again Case 1, with a match length of 4 — the match is `ATTA` — so we backward step and find 3 copies of $ ≠ *P* [4] = `A` in rows 30–32.

This is now Case 3 with a match length of 5, so we pass to the LCE data structure the SA values for rows 30 and 32 (which we know by decrementing when we backward step) and the sampled SA entries for the close 3-subsequences of the previous occurrence BWT[28] and next occurrence BWT[33] of *P* [4] = `A`. The LCE data structure tells us to continue with the 3 copies of `A` in rows 26– 28, now with a match length of only 3 — the match is `GAT`. Because the match length has decreased, we report that the last 3-MEM of *P* with respect to *T* is `GATTA`.

We are now in Case 1 again, so we backward step and find 2 copies of *P* [3] = `T` and 1 copy of `$`, in rows 10–12. This is Case 2 with a match length of 4 — the match is `AGAT` — so we change the range to the close 3-subsequence 9–11 of the *P* [3] = `T` in row 11 and set the match length to the minimum of 4 and the length 1 of the longest common prefix of the corresponding suffixes. Because the match length has decreased, we report that the penultimate 3-MEM of *P* with respect to *T* is `AGAT`.

We backward step and find 3 copies of `T` in rows 40–42, with match length 2 for `TA`. Because *P* [2] = `A`, we are in Case 3, so we again use the SA values and the LCE data structure to decide when to continue with the 3 copies of `A` in rows 37–39, or the ones in rows 43–45. The LCE data structure tells us to continue with the former, with a match length of 2 for `TA`.

We backward step and find 3 copies of *G* in rows 17–19, with match length 3 for `ATA`. Because *P* [1] = `C`, we are in Case 3 again. The SA entries and LCE data structure tell use to continue with the close 3-subsequence of the `C` in row 15, with match length 2 for `AT`. Because the match length has decreased, we report that `ATA` is another 3-MEM of *P* with respect to *T*.

Finally, since we have reached the left end of *P*, we report that `CAT` is a 3-MEM of *P* with respect to *T*.

## 4 Conclusion

We have defined *k*-MEMs as a generalization of MEMs, shown how MONI can easily be extended to MONI-*k* to find them efficiently, and proposed how they can be applied in pangenome-to-pangenome comparison. We leave as future work developing a similar data structure that takes *k* at query time, as well as extending the implementation of MONI to obtain an actual implementation of MONI-*k*, and applying it in bioinformatics research.

## Footnotes

1 We also let *T* be a set of strings with total length *n* instead of a single string. We note as an aside that in the stringology community MEMs are sometimes called super-maximal exact matches, with “MEM” meaning something slightly different.

2 A recent tool by Arakawa et al. [1], called a bidirectional r-index, can handle many genomes but cannot delete and so apparently cannot be used to find MEMs efficiently.

3 By “pangenome”, here we mean simply a sufficiently large collection of genomes that most reasonably common variants are likely to be represented.

4 We refer readers unfamiliar with *r* to Kempa and Kociumaka’s recent paper [8], in which they showed *r* is never more than an *O*(log^2^ *n*)-factor larger than the number of phrases in the LZ77 parse of *T* — which is very small when *T* is a highly repetitive dataset such as a pangenome.

5 Although we cannot show the SLP takes less space than the r-index in the worst case, it does in practice.

6 We note that it is easy to build an index with which we can find *k*-mers that are fairly common in *T*_1_ but fairly rare in *T*_2_, for some *k* given at construction time, but this is incomparable with what we are doing.

7 The reader may notice we do not need the first and last rows; we leave them in for consistency.

